# Distinct disease-modifying therapies are associated with different blood immune cell profiles in people with relapsing-remitting multiple sclerosis

**DOI:** 10.1101/2023.07.20.549852

**Authors:** João Canto-Gomes, Daniela Boleixa, Catarina Teixeira, Ana Martins da Silva, Inés González-Suárez, João Cerqueira, Margarida Correia-Neves, Claudia Nobrega

## Abstract

Disease modifying therapies (DMTs) used for treating people with relapsing-remitting multiple sclerosis (pwRRMS) target the immune system by different mechanisms of action. However, there is a lack of a comprehensive comparison of their effects on the immune system. Herein, we evaluated the numbers of circulating B cells, CD4^+^ and CD8^+^ T cells, regulatory T cells (Tregs), natural killer (NK) cells and NKT cells, and their subsets, in pwRRMS who were treatment-naïve or treated with different DMTs. Compared to treatment-naïve pwRRMS, common and divergent effects on immune system cells were observed on pwRRMS treated with different DMTs, with no consistent pattern across all therapies in any of the cell populations analysed. PwRRMS treated with fingolimod, dimethyl fumarate (DMF), or alemtuzumab have reduced numbers of CD4^+^ and CD8^+^ T cells, as well as Treg subsets, with fingolimod causing the most pronounced decrease in T cell subsets. In contrast, teriflunomide and interferon (IFN) β have minimal impact on T cells, and natalizumab marginally increases the number of memory T cells in the blood. The effect of DMTs on the B cell, NKT and NK cell subsets is highly variable with alemtuzumab inducing a strong increase in the number of the most immature NK cells and its subsets. This study highlights the absence of a consistent pattern of the impact of various DMTs on immune system cells, with variations in both direction and magnitude of effect thus reenforcing the notion that distinct immune cell subsets are potential players in MS pathophysiology and/or DMT efficacy.

## 1. Introduction

Multiple sclerosis (MS) is an autoimmune disease of the central nervous system (CNS) that affects more than 2.8 million people worldwide (1). Most people with MS (pwMS) are diagnosed with relapsing-remitting MS (RRMS; ∼85%), which is characterised by interleaved episodes of disability and recovery (2). There is no cure for MS, but several therapies have been shown to be effective in preventing disease progression in pwRRMS (3-5).

Most therapies available for RRMS dampen the immune response, either by preventing cells from entering the CNS, by lowering specific immune cell activities (including cell polarisation, proliferation, and peptide presentation) or even by inducing immune cell depletion. Several disease-modifying therapies (DMT) are available for MS and the decision supporting the choice of the appropriate DMT for a given person is complex and mostly based on the individual clinical profile (*e.g*., disease activity, safety, comorbidities, side effects and pregnancy planning). DMTs can be categorised according to their efficacy: modest, moderate, and high. The increasing efficacy of DMTs correlates with increasing side effects and complications, including the risk of lymphopenia, infection, development of secondary autoimmune disorders and development of progressive multifocal leukoencephalopathy (6-9). In severe cases of MS, an intensive therapeutic approach using high-efficacy DMTs is preferred to reduce progression. In pwRRMS with lower disease activity, DMTs with more modest efficacy are chosen initially and escalated to more effective DMTs when needed. DMTs with modest efficacy include glatiramer acetate, interferon (IFN)β and teriflunomide; moderate to high efficacy include fumarate derivatives [e.g., dimethyl fumarate (DMF)], cladribine and sphingosine-1-phosphate (S1P) inhibitors (e.g., fingolimod); and high efficacy include alemtuzumab, mitoxantrone, natalizumab, anti-CD20 monoclonal antibodies, and autologous haematopoietic stem cell transplantation (10, 11).

Existing studies evaluating the effects of different DMTs on pwRRMS mostly focus on their efficacy in preventing relapses, formation of new lesions and disease progression, providing very scarce data on the effects of different DMTs on immune cell subsets and phenotypes. Although the mechanism of action of some DMTs is partially understood, there are few studies with extensive phenotypic characterisation of immune system cells. Indeed, most studies have focused on the effects of therapies on conventional T and, more recently, on B cells, which are considered the most relevant immune cell subsets involved in the disease (4). However, there is increasing evidence supporting the notion that several other immune system cells with regulatory potential are involved in MS, including regulatory T cells (Tregs), natural killer (NK) cells and NKT cells. Tregs have an essential role in preventing autoimmunity and the percentage of these cells has been found to increase with therapy, including IFNβ (12). NK and NKT cells may have important implications in MS as they are able to produce both pro- and anti-inflammatory cytokines, depending on the local cytokine *milieu*, and can mediate cellular cytotoxicity against abnormal cells through the engagement of their inhibitory and activating receptors (13-16). Interestingly, treatment with some DMTs, including natalizumab and alemtuzumab, has been shown to be associated with an increased frequency and number of CD56^bright^ NK cells (17). However, data on the phenotype of NK and NKT cell subsets, including the expression of inhibitory (*e.g*., KIR3DL1, KIR2DL2/3, KLRG1 and NKG2A) and activating (*e.g*., NKp30, NKp44 and NKp46) receptors, in the context of different MS therapies remains scarce.

Comparative studies on the impact of different DMTs on immune system cells are important to better understand whether the degree of efficacy of DMTs correlates with the magnitude of their impact on immune cell subsets and whether there is a common immune target between the different DMTs. The aim of this study was to compare the effects of different DMTs on blood immune cell subsets. For this purpose, a comprehensive phenotypic characterization of CD4^+^ and CD8^+^ T cells, Tregs, NK and NKT cells was performed in treatment-naïve pwRRMS and compared with pwRRMS treated with one of the following DMTs: alemtuzumab, DMF, fingolimod, IFNβ, natalizumab or teriflunomide.

## 2. Material and Methods

### 2.1. Participants

This multicentre cross-sectional study was approved by the local Ethical Committees and participants were recruited at the Hospital de Braga (ref. 5888/2016-CESHB; Braga, Portugal), Hospital Geral de Santo António (ref. 098-DEFI/097-CES; Porto, Portugal) and Hospital Alvaro Cunqueiro (ref. 2021/430; Vigo, Spain) between 2018 and 2022. The inclusion criteria were: diagnosis of RRMS according to McDonald 2017 criteria, over 18 years of age and in the same DMT for more than 6 months prior to recruitment. Excluding criteria were: other immune-mediated diseases in addition to MS; history of treatment for cancer; splenectomy or thymectomy; pregnancy; treatment with corticosteroids in the previous 3 months or continuously for more than 6 months. Participants signed a written informed consent and agreed to provide demographic and clinical data, obtained from their medical records, including age, sex, age of MS onset, months living with MS, time since last DMT administration, and disease severity [Expanded Disability Status Scale (EDSS); **Table 1**]. The control group of recently diagnosed treatment-naïve pwRRMS with a diagnosis of RRMS according to McDonald 2017 criteria, aged over 18 old, and not meeting any of the above exclusion criteria has been described previously (18).

**Table 1.**
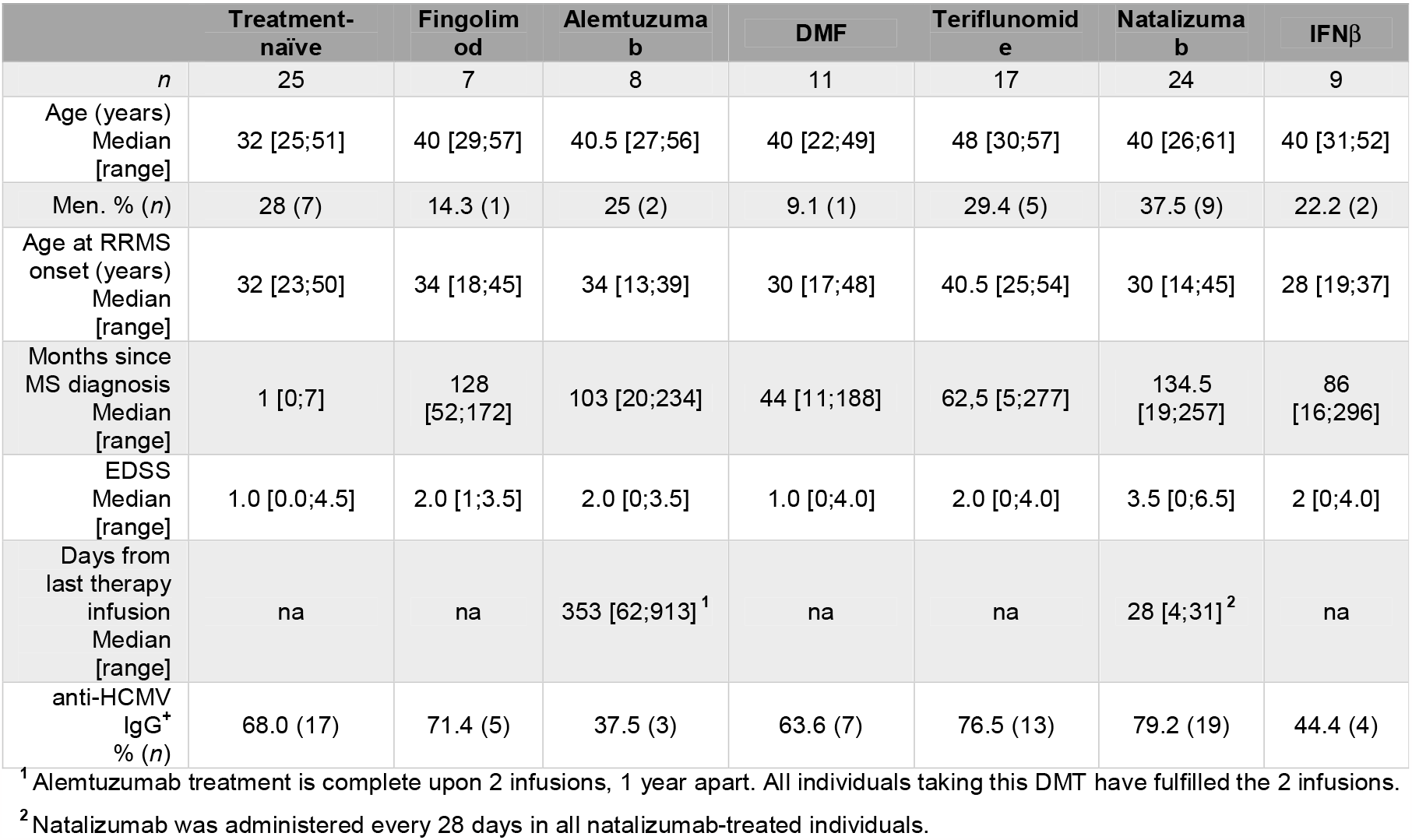
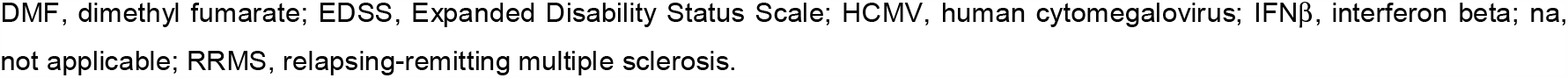
Clinical and demographic data of the study groups.

|  | Treatment-naïve | Fingolimod | Alemtuzumab | DMF | Teriflunomide | Natalizumab | IFNβ |
| --- | --- | --- | --- | --- | --- | --- | --- |
| *n* | 25 | 7 | 8 | 11 | 17 | 24 | 9 |
| Age (years)<br>Median<br>[range] | 32 [25;51] | 40 [29;57] | 40.5 [27;56] | 40 [22;49] | 48 [30;57] | 40 [26;61] | 40 [31;52] |
| Men. % (*n*) | 28 (7) | 14.3 (1) | 25 (2) | 9.1 (1) | 29.4 (5) | 37.5 (9) | 22.2 (2) |
| Age at RRMS onset (years)<br>Median<br>[range] | 32 [23;50] | 34 [18;45] | 34 [13;39] | 30 [17;48] | 40.5 [25;54] | 30 [14;45] | 28 [19;37] |
| Months since MS diagnosis<br>Median<br>[range] | 1 [0;7] | 128 [52;172] | 103 [20;234] | 44 [11;188] | 62.5 [5;277] | 134.5 [19;257] | 86 [16;296] |
| EDSS<br>Median<br>[range] | 1.0 [0.0;4.5] | 2.0 [1;3.5] | 2.0 [0;3.5] | 1.0 [0;4.0] | 2.0 [0;4.0] | 3.5 [0;6.5] | 2 [0;4.0] |
| Days from last therapy infusion<br>Median<br>[range] | na | na | 353 [62;913]<sup>1</sup> | na | na | 28 [4;31]<sup>2</sup> | na |
| anti-HCMV IgG+<br>% (*n*) | 68.0 (17) | 71.4 (5) | 37.5 (3) | 63.6 (7) | 76.5 (13) | 79.2 (19) | 44.4 (4) |

<sup>1</sup> Alemtuzumab treatment is complete upon 2 infusions, 1 year apart. All individuals taking this DMT have fulfilled the 2 infusions.
<sup>2</sup> Natalizumab was administered every 28 days in all natalizumab-treated individuals.
DMF, dimethyl fumarate; EDSS, Expanded Disability Status Scale; HCMV, human cytomegalovirus; IFNβ , interferon beta; na, not applicable; RRMS, relapsing-remitting multiple sclerosis.

### 2.2. Sample Processing

Peripheral blood processing, absolute cell numbers quantification, human cytomegalovirus (HCMV) serology and FACS staining were performed as previously reported (18, 19). For FACS staining, per participant, a vial containing 5 million PBMCs was thawed and divided by four distinct panels of antibodies (**Supplementary Table 1**). The phenotype of the different immune cell subsets evaluated is shown in **Supplementary Table 2**.

### 2.3. Statistical Analysis

The sample size and the median number of each cell population studied per each study group are summarized in **Supplementary Table 3**. The contribution of each DMT, controlling for age, sex and HCMV serology, on the number of every immune cell subset, was evaluated through a two-stage multiple linear regression analysis, as previously reported (20). In the first regression model, the number of every immune cell subset (Log_10_-transformed to overcome the non-normal distribution, determined by the D’Agostino & Pearson test) was included as the dependent variable; age, sex and HCMV serology were included as independent variables. Outputs from these regressions are depicted in **Supplementary Table 4**. The unstandardized residuals from each regression were retrieved to be used in a second regression model; these unstandardized residuals of a regression model represent the variation on the dependent variable (*i.e*., the number of every immune subset) that is not explained by the independent variables (*i.e*., age, sex and HCMV serology). In the second regression, the unstandardized residuals were used as the dependent variable and the different DMTs as the independent variables. Therapies were included in the regression as dummy variables having as reference the treatment-naïve pwRRMS. Statistical outputs of these regressions are shown in **Supplementary Table 5**.

In the case of natalizumab- or alemtuzumab-treated pwRRMS, the impact of the time since the last drug infusion on the immune cell subsets’ numbers was evaluated by performing regression models using the unstandardized residuals from the abovementioned first regressions (dependent variable) and time since last drug infusion (independent variable). These regression outputs are represented in **Supplementary Tables 6** and **7**.

All regression analyses were performed using IBM SPSS Statistics v26 (IBM Corporation, NY, USA); GraphPad Prism v9 (GraphPad Software, CA, USA) was used for the graphical representation of the heatmaps. Differences were considered statistically significant when the *p*-value <0.050. In the regression models, the magnitude of the effect of each independent variable was inferred based on the absolute standardized *β*-values and classified as: small, when |*β*-value| <0.29; medium, when 0.30< |*β*-value| <0.49, or large, when |*β*-value| >0.5 (21). During flow cytometry analysis, cell subsets whose parent population was represented with less that 500 events were not considered for the statistical analysis. For this reason, the number of individuals analysed varied between cell subsets of the same group.

## 3. Results

### 3.1. Fingolimod, alemtuzumab and DMF have a strong negative effect on T cells, as opposed to teriflunomide, natalizumab and IFNβ

The blood absolute numbers of different CD4^+^ and CD8^+^ T cell subsets, Tregs, NK and NKT cells in pwRRMS treated with one out of the six distinct DMTs compared to treatment-naïve pwRRMS were assessed. Fingolimod- and DMF-treated pwRRMS had lower total lymphocyte counts compared to treatment-naïve pwRRMS (**Figure 1**). In contrast, individuals treated with natalizumab presented a slight increase in total lymphocyte counts. The effect of DMTs on the number of B cells was quite heterogeneous across DMTs, with alemtuzumab and natalizumab leading to higher numbers of B cells compared to treatment-naive pwRRMS, whereas fingolimod induced lower numbers of those cells. Regarding the T cell subsets, fingolimod, alemtuzumab or DMF treatment led to a significant reduction in the number of most subsets, with fingolimod affecting the naive CD4^+^ and CD8^+^ T cells strongly, as opposed to alemtuzumab and DMF that induced a stronger effect in the memory T cell pool. Natalizumab had an effect in increasing total T cell numbers, specifically of memory subsets. Teriflunomide and IFNβ had a residual impact on T cell subsets.

**Figure 1.**
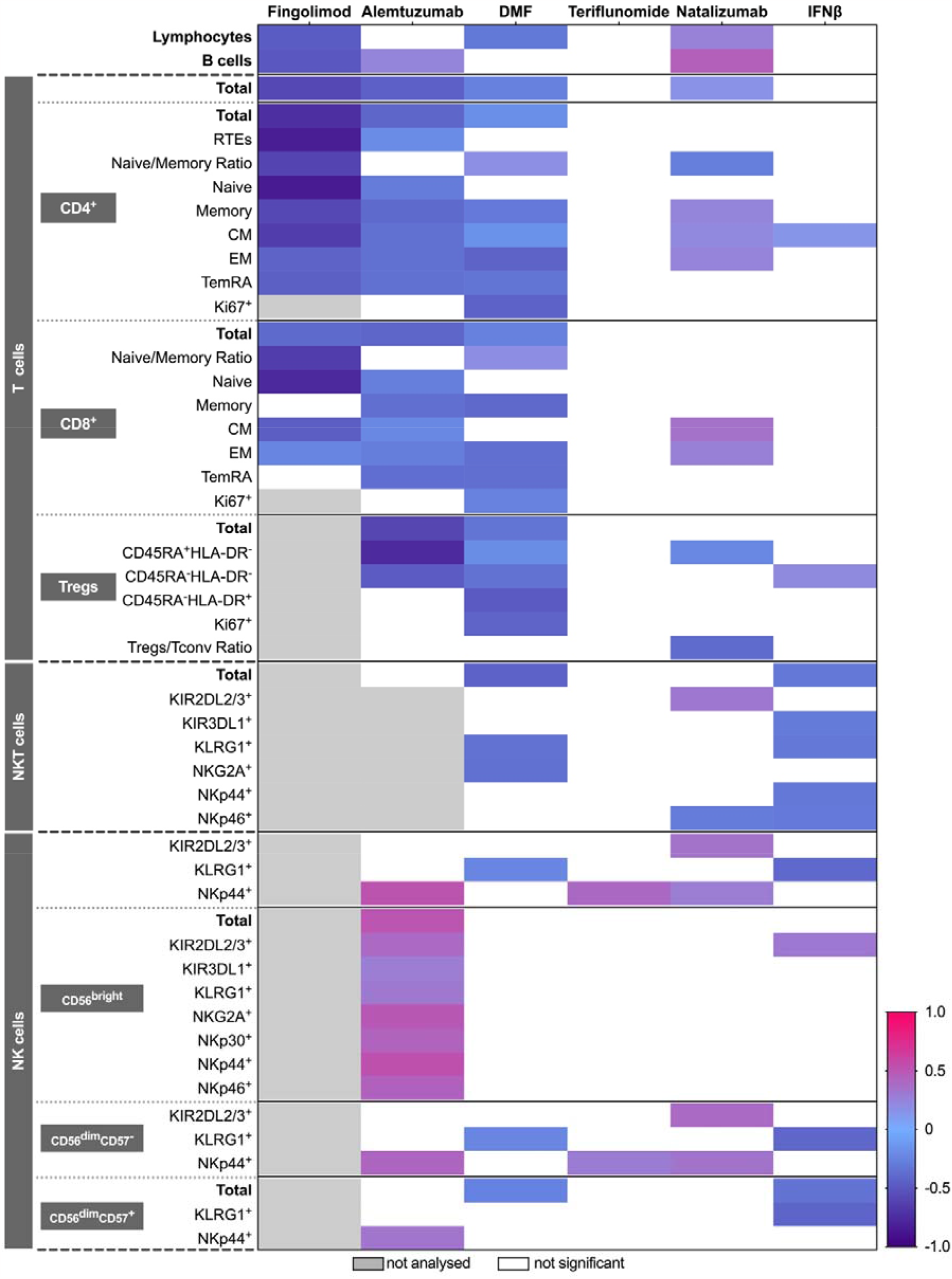
Effect of multiple sclerosis (MS) disease-modifying therapies (DMTs) on the number of immune cell subsets. The effect of DMTs on the number of immune cell populations was assessed using sequential multiple linear regression models. To exclude the effect of age, sex, and human cytomegalovirus (HCMV) serology on immune cell counts, a first multiple linear regression was performed for each cell subset (**Supplementary Table 4**). A second regression was performed using the residuals of the first regression (the part not explained by age, sex and HCMV serology) as the dependent variable and the DMTs (alemtuzumab, DMF, fingolimod, IFNβ, natalizumab or teriflunomide; using treatment-naïve pwRRMS as reference) as the independent variable. For each cell subset, standardised β-values of the second regression were obtained (**Supplementary Table 5**), reflecting the strength of the effect of each treatment (independent variable) on the number of the cell subset (dependent variable) compared with treatment-naïve pwRRMS. These standardised β-values were used to generate a heatmap of the effect of the different DMTs on the number of immune cells. On the heatmap, grey cells represent cell subsets that were not analysed; white cells represent predictors that were not significant in the regression models. Only regression models with at least one statistically significant standardised β-values are shown on the heatmap. CM, central memory; EM, effector memory; DMF, dimethyl fumarate; IFNβ, interferon beta, RTE, recent thymic emigrants; Tconv, conventional CD4^+^ T cells; TemRA, terminally differentiated effector memory T cells re-expressing CD45RA; Treg, regulatory T cells; NK, natural killer cells.

### 3.2. DMTs mostly led to decreased numbers of NKT cells, with no clear signature on NK cells distinguishing the treated pwRRMS

Overall, pwRRMS receiving different DMTs were found to present reduced numbers of distinct NKT cell subsets (**Figure 1**). The small number of pwRRMS with sufficient cells for flow cytometry analysis limited the evaluation of NKT cells in individuals receiving fingolimod (n=2) or alemtuzumab (n=2), and for this reason these groups were excluded from the analysis of NKT cells. Both DMF- and IFNβ-treated pwRRMS showed a decrease in total NKT cell numbers though having impact in different subsets: DMF induced a great reduction in the number of NKT cells expressing KLRG1 and NKG2A whereas IFNβ impacted the most those expressing KIR3DL1, KLRG1, NKp44 and NKp46 receptors. Teriflunomide had no effect on NKT cell numbers and its subsets, while natalizumab caused an increase in the number of KIR2DL2/3^+^ NKT cells and a decrease in NKp46 NKT cells.

The effect of different DMTs on NK cell subset counts varied greatly in comparison to treatment-naïve pwRRMS. Alemtuzumab-treated pwRRMS had increased numbers of NKp44^+^ on all NK cell subsets. Additionally, the numbers of CD56^bright^ NK cell expressing any of the activation and inhibitory receptors analysed were increased in alemtuzumab-treated pwRRMS. In contrast, DMF treatment leads to a reduction in the number of total CD56^dim^CD57^+^ NK cells, and in the number of total and CD56^dim^CD57^-^ NK cells expressing KLRG1. Teriflunomide treatment resulted in increased numbers of total and CD56^dim^CD57^-^ NK cells expressing NKp44. Compared to treatment-naïve individuals, natalizumab-treated pwRRMS had higher numbers of KIR2DL2/3^+^ and/or NKp44^+^ cells among total and CD56^dim^CD57^-^ NK cells. IFNβ-treated pwRRMS had higher counts of KIR2DL2/3^+^ CD56^bright^ NK cells, and lower counts of CD56^dim^CD57^+^ NK cells and total and of both CD56^dim^ NK cell subsets expressing KLRG1.

### 3.3. Time since the last infusion of alemtuzumab or natalizumab correlates with changes in the number of immune cell subsets

The therapeutic regimen of alemtuzumab and natalizumab involves intervals between successive drug infusions, as opposed to the other studied DMTs that are administered daily (fingolimod, teriflunomide, DMF) or every other day (IFNβ). As pwRRMS treated with alemtuzumab and natalizumab differ greatly in the time since the last drug administration (**Table 1**), we performed a regression analysis to evaluate the relationship between the time since the last drug infusion and the number of circulating immune cells (**Figure 2**).

**Figure 2.**
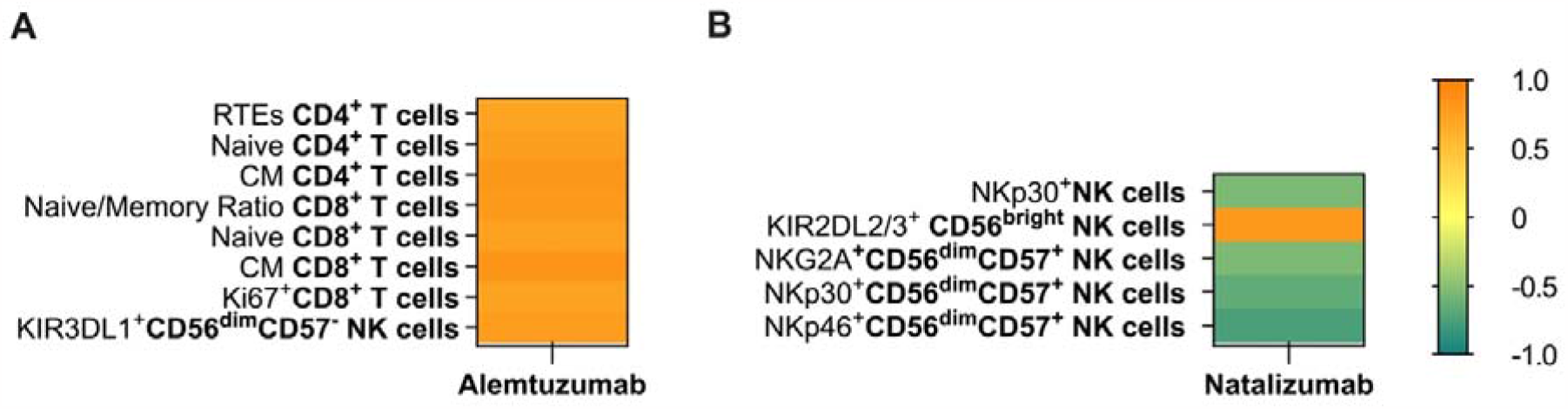
Effect of time since last alemtuzumab or natalizumab infusion on the number of immune cell subsets. The impact of time since the last alemtuzumab (**A**) or natalizumab (**B**) infusion on the number of immune cell populations was assessed using sequential multiple linear regression models. The effect of age, sex, and human cytomegalovirus (HCMV) serology on immune cell counts was assessed using a first multiple linear regression. Using the residuals from the first regression (the part not explained by age, sex and HCMV serology), a second regression was performed for each cell subset with these residuals as the dependent variable, and time since the last drug infusion as the independent variable. From these regressions, standardised β-values were obtained and used to generate a heatmap of the effect of time since the last alemtuzumab (**Supplementary Table 6)** or natalizumab infusion (**Supplementary Table 7)** on the number of immune cell subsets. Only significant linear regression models are shown on the heatmaps. CM, central memory; NK, natural killer cells; RTEs, recent thymic emigrants.

Increasing time since the alemtuzumab infusion were related with higher numbers of RTEs CD4^+^ T cells and of naïve and CM among CD4^+^ and CD8^+^ T cells, which in the case of CD8^+^ T cells resulted in a higher ratio of naïve to memory cells. In addition, higher numbers of proliferating CD8^+^ T cells and of KIR3DL1^+^ CD56^dim^CD57^-^ NK cells were observed with increasing time since the last alemtuzumab infusion.

The effect of time since the last natalizumab infusion in pwRRMS was observed only on NK cells. A negative association was observed between the time since the last natalizumab infusion and the number of NKp30^+^ NK cells, and of CD56^dim^CD57^+^ NK cells expressing NKG2A, NKp30 and/or NKp46. In contrast, the number of KIR2DL2/3^+^ CD56^bright^ NK cells increased with time since the last natalizumab infusion.

Overall, while the time since the last alemtuzumab infusion correlated mostly with changes in the number of T cell subsets, the time since the last natalizumab infusion correlated with changes in NK cells.

## 4. Discussion

We provide a comprehensive comparison of the effect of different DMTs on the number of blood immune cell populations in pwRRMS. Control for age, sex and HCMV serology was essential to generate robust information, as these variables are known to influence immune cell homeostasis (22, 23). The analysis performed in this study allowed to dissect the direction and magnitude of the impact of different DMTs used for MS on immune system cells. PwRRMS treated with DMF, alemtuzumab or fingolimod presented a strong decrease in the number of most T cell subsets. Conversely, pwRRMS treated with teriflunomide or IFNβ presented virtually no alterations in T cells numbers while those treated with natalizumab showed a modest increase in T cell subsets’ numbers in blood. There was no clear pattern in the effect of DMTs on the number of the other analysed immune system cells (NKT and NK cells).

The effect of fingolimod, DMF or alemtuzumab in T cell subsets is consistent with the proposed mechanism of action of these DMTs. Fingolimod acts as a functional antagonist of S1P receptors, which are essential for lymphocyte egress from the lymph nodes, overriding the CCR7 retention signal. By blocking S1P receptors, CCR7-expressing cells (*i.e*., naive and CM T cells) are retained in the lymph nodes (24). Decreased percentages and numbers of naïve and CM T cells have been described in the blood of fingolimod-treated pwRRMS compared to treatment-naïve pwRRMS (24, 25). In addition, Ghadiri *et al*. also showed that the numbers of RTEs and EM T cells are decreased in fingolimod-treated individuals, which correlates with our observations (25). Though fingolimod acts more strongly on RTEs, naïve and CM T cells, the cumulative sequestration of those cells in lymph nodes might contribute to the reduced number of the cells that succeed in the differentiation process, namely the EM T cells. Regarding DMF, despite its direct antioxidant effects on the nervous system and neuroprotection, it has been described to induce T cell apoptosis, particularly of those with a memory phenotype (EM and TemRA), and reduce their activation and proliferation (26, 27). Consistent with this, we found that DMF-treated pwRRMS showed a reduction in memory, but not in naive T cell subsets, and decreased numbers of proliferating T cells. Alemtuzumab is described to lead to targeted cytotoxicity of all CD52-expressing cells (*i.e*., T and B cells, and monocytes). Baker *et al*. observed that the numbers of CD4^+^ and CD8^+^ T cells, and of Tregs recovered to only ∼30% to ∼50% of the baseline levels after one year of alemtuzumab infusion (9). Moreover, Haas *et al*. showed a sustained increase in the percentage of CD45RO^+^ Tregs (28). We now add that the number of naïve Tregs (CD45RA^+^HLA-DR^-^) is reduced to a greater extent in comparison to memory (CD45RA^-^) Tregs, in alemtuzumab-treated compared to treatment-naïve pwRRMS. Given the broad effect of alemtuzumab on all T cell subsets, more immune cell subsets were expected to correlate with time since the last alemtuzumab infusion. A possible explanation is that those immune cell subsets whose numbers recover rapidly after drug infusion, may not have been captured in the regression models due to the wide variation of the time elapsed since the last alemtuzumab infusion of our cohort.

There was no clear pattern on T cell subsets on the effect of teriflunomide and IFNβ, having little to no effect. Teriflunomide inhibits the dihydroorotate dehydrogenase and consequently the *de novo* pyrimidine synthesis, thus limiting cellular proliferation, particularly of activated T cells (29, 30). Despite this, no effect of teriflunomide was observed on the number of proliferating CD4^+^ or CD8^+^ T cells. The IFNβ is described to induce T cell polarisation towards a Th2 phenotype, to the detriment of Th1 or Th17 (31, 32). We now added that IFNβ does not lead to major alterations in T cell subset numbers. Regarding natalizumab, by binding the α-4 integrin of VLA-4 on lymphocytes, it inhibits cell migration to the CNS and leads to their accumulation in the peripheral blood (33-35). We showed an increase mainly in the numbers of memory T cell subsets in natalizumab-treated pwRRMS, compared to treatment-naïve. Picker *et al*. suggested that the accumulation of memory T cells in the periphery, but not of naïve T cells, may be related to the higher expression of the α-4 integrin on memory subsets (36). The greater increase in peripheral CM CD8^+^ T cells in comparison to other CD8^+^ or CD4^+^ memory T cell subsets showed by us may correlate with the fact that CM T cells predominate in the CSF and that CD8^+^ T cells are the most abundant T cells in MS lesions (37).

We found that pwRRMS treated with fingolimod had reduced numbers of B cells. Fingolimod prevents B cells migration from lymphoid organs, leading to their reduction in the peripheral blood. In contrast, alemtuzumab and natalizumab treatment increased the numbers of B cells in the peripheral blood. While in natalizumab-treated pwRRMS the higher number of B cells in the peripheral blood may be due to blockage of cell migration into the tissues, in alemtuzumab-treated pwRRMS the higher number of B cells is intriguing and seems to contradict the depleting effect of the therapy. However, Baker *et al*. observed that although alemtuzumab treatment leads to an immediate and dramatic reduction in B cell counts, it is followed by a rapid hyper-repopulation to values exceeding baseline levels one year after drug infusion (9). In line with this report, here the alemtuzumab-treated group of pwRRMS had a median time since the last drug infusion of nearly 12 months. The rapid repopulation of B cells together with the lower Treg counts in alemtuzumab-treated pwRRMS may be a relevant finding considering that this drug predisposes to B cell-mediated secondary autoimmune diseases (7).

In treated pwRRMS, no apparent pattern of DMT’s effect on NK and NKT cell subset numbers could be seen. The numbers of distinct NKT cell subsets were reduced in response to DMF and IFNβ. Others have previously found fewer NKT cells in DMF-treated pwRRMS, but the impact of DMF on the underlying NKT cell subsets was not examined (27, 38). In IFNβ- or DMF-treated pwRRMS, fewer KLRG1^+^ NKT cells were observed. KLRG1^+^ NKT cells have been found to be long-lived effector cells that can react to antigens (namely, α-Galactosylceramide), according to data from mice (39). Further research is necessary to determine whether these NKT cells have a role in the pathogenesis of MS in humans, and whether their decline is connected to the effectiveness of DMF and IFNβ in the treatment of pwRRMS.

Regarding the DMTs effect on NK cells, we found that alemtuzumab, teriflunomide and natalizumab resulted in increased numbers of NKp44^+^ NK cells, suggesting that these cells may play a role in the mechanism of action or efficacy of these DMTs. Our study supports previous findings showing increased CD56^bright^ NK cells in alemtuzumab-treated pwRRMS (40, 41). The expression of VLA-4 by NK cells provides a plausible explanation for the increased numbers of NK cell subsets in circulation during natalizumab treatment (42). Notably, the time since the last natalizumab infusion negatively correlated with NK cell subset counts. Nevertheless, it remains unknown whether this decline is associated with the recovery of migratory potential and with the migration of these cells to tissues.

DMTs described as less effective (*e.g*., teriflunomide and IFNβ) are the ones with the least effect on the number of T cell subsets. Nevertheless, conclusions from these observations should be drawn with caution as these DMTs are used in pwRRMS with slower and less severe disease progression, as opposed to DMTs of higher efficacy (*e.g*., alemtuzumab, natalizumab) (10, 11). It is not known whether the effect of high *vs*. modest efficacy DMTs is an exclusive consequence of the DMT itself and/or a consequence of intrinsic changes in immune cell alterations that predispose to distinct disease severity and consequently the need for high/modest efficacy DMTs.

Overall, we observed moderate to strong alterations in the number of blood immune cells in pwRRMS in all DMTs evaluated, including in those groups with small sample sizes, namely pwRRMS treated with fingolimod or alemtuzumab (n=7 and n=8, respectively). Although the differences observed in the number of T cell subsets correlate with the expected and described mechanism of action of different DMTs, this study demonstrates that the magnitude of the impact of the various MS therapies on immune cells is different, namely on T cells. Additionally, it underlines the contrasting impact of distinct DMTs on subsets of NK and NKT cells, further reinforcing their potential involvement in MS pathophysiology and/or DMT efficacy.

## Data availability statement

The data that support the findings of this study are available from the corresponding author upon request.

## Author Contributions

JC-G, MC-N, JJC, and CN contributed to conception and design of the study. JJC, AMdS, DB, CT and IG-S included patients and provided clinical data. JC-G and CN organized the database; performed the experimental tasks and statistical analysis; generated the figures and tables; and wrote the first draft. All authors contributed to the article, including to manuscript revision, and approved the submitted version.

## Acknowledgments

The authors would like to thank the people with multiple sclerosis that agreed to participate in this study.

